# The mammalian Ire1 inhibitor, 4µ8C, exhibits broad anti-*Aspergillus* activity *in vitro* and in a treatment model of fungal keratitis

**DOI:** 10.1101/2024.08.08.607189

**Authors:** Manali M. Kamath, Emily M. Adams, Jorge D. Lightfoot, Becca L. Wells, Kevin K. Fuller

## Abstract

**Objective:** The fungal unfolded protein response consists of a two-component relay in which the ER-bound sensor, IreA, splices and activates the mRNA of the transcription factor, HacA. Previously, we demonstrated that *hacA* is essential for *Aspergillus fumigatus* virulence in a murine model of fungal keratitis (FK), suggesting the pathway could serve as a therapeutic target. Here we investigate the antifungal properties of known inhibitors of the mammalian Ire1 protein both *in vitro* and in a treatment model of FK.

**Methods:** The antifungal activity of Ire1 inhibitors was tested against conidia of several *A. fumigatus* isolates by a microbroth dilution assay and against fungal biofilm by XTT reduction. The influence of 4μ8C on *hacA* mRNA splicing in *A. fumigatus* was assessed through gel electrophoresis and qRT-PCR of UPR regulatory genes. The toxicity and antifungal profile of 4μ8C in the cornea was assessed by applying drops to uninfected or *A. fumigatus*-infected corneas 3 times daily starting 4 hours post-inoculation. Corneas were evaluated daily through slit-lamp imaging and optical coherence tomography, or at endpoint through histology or fungal burden quantification via colony forming units.

**Results:** Among six Ire1 inhibitors screened, the endonuclease inhibitor 4μ8C displayed the strongest antifungal profile with an apparent fungicidal action. The compound both blocked conidial germination and hyphal metabolism of *A. fumigatus* Af293 in the same concentration range that blocked *hacA* splicing and UPR gene induction (60-120 µM). Topical treatment of sham-inoculated corneas with 0.5 and 2.5 mM 4μ8C did not impact corneal clarity, but did transiently inhibit epithelialization of corneal ulcers. Relative to vehicle-treated Af293-infected corneas, treatment with 0.5 and 2.5 mM drug resulted in a 50% and >90% reduction in fungal load, respectively, the latter of which corresponded to an absence of clinical signs of infection or corneal pathology.

**Conclusion:** The *in vitro* data suggest that 4μ8C displays antifungal activity against *A. fumigatus* through the specific inhibition of IreA. Topical application of the compound to the murine cornea can furthermore block the establishment of infection, suggesting this class of drugs can be developed as novel antifungals that improve visual outcomes in FK patients.

## 1 Introduction

Fungal infections of the cornea are an ophthalmologic emergency and represent a leading cause of ocular morbidity and unilateral blindness worldwide (1,2). This disease entity, called fungal keratitis (FK), typically affects otherwise healthy individuals that are inoculated with fungal spores or hyphae through ocular trauma or contact lens wear (3,4). Even with the current standard of treatment, which includes topical application of natamycin or voriconazole, 40-60% of all FK cases result in corneal perforation and/or the need for corneal transplantation (5,6). One approach to the development of novel antifungals for FK, or indeed any clinical context, is a biologically-informed one that involves two key steps: first, the identification of a fungal protein/pathway that is essential for growth or virulence in a relevant disease model and, second, identification of a small molecule(s) that inhibits protein function and improves disease outcome in the model without toxicity to the host. In the context of this study, we and others have previously identified the unfolded protein response (UPR) as a critical regulator of *Aspergillus fumigatus* virulence in both lung and cornea infection models, and now seek to develop inhibitors of the pathway as novel antifungals (7–9).

The mammalian UPR consists of three separate but partially redundant signaling branches – IRE1, ATF6, and PERK-where the Ire1 branch is the most highly conserved across the eukaryotes and is the only one found in the fungi (10–14). In *A. fumigatus*, the Ire1 ortholog (IreA) is an ER-bound transmembrane protein comprised of a luminal sensory domain and two cytosolic domains that propagate the signal to the nucleus (7,15). More completely, the rapid accumulation of misfolded proteins in the ER lumen promotes the oligomerization of IreA within the membrane. This clustering triggers the trans-autophosphorylation of the kinase domain and subsequent activation of the endoribonuclease domain which then splices an unconventional intron from its only known client mRNA, *hacA*. The spliced *hacA* isoform encodes a bZIP transcription factor, HacA, which regulates genes not only involved directly in protein folding homeostasis (e.g. chaperones and foldases), but also genes involved in primary and secondary metabolism that altogether promote adaptation to stressful environments (7,9,16,17). Recently, we demonstrated that a *hacA* deletion mutant of *A. fumigatus* (Af293 background) is viable under normal growth conditions but is unable to establish infection in a murine model of fungal keratitis (9). Surprisingly, we were unable to isolate an *ireA* deletant in Af293 and repression of the gene using a tetracycline regulatable promoter blocks hyphal growth (9). Not only does this indicate that IreA regulates fungal growth beyond its influence of HacA, but it pragmatically suggests the protein could serve as an ideal target for antifungal intervention.

Several small molecules are known to inhibit the mammalian Ire1 kinase or endonuclease domains and, consistent with the UPR’s role in regulating cellular homeostasis, display anti-proliferative effects various cell culture or *in vivo* cancer models, including those related to hepatocellular carcinoma, acute myeloid leukemia, triple negative breast cancer, and multiple myeloma (15–21). Recently, Guirao-Abad and colleagues demonstrated that one such compound, the coumarin derivative 4-methyl umbilliferone 8-carbaldehyde (4µ8C), inhibits the endonuclease activity of *A. fumigatus* IreA and consequently the downstream expression of canonical HacA-dependent genes in the setting of acute ER stress (18). The utility of 4µ8C or other Ire1 inhibitors as clinically useful antifungals, however, has not been explored. In this study, we assess the intrinsic antifungal activity of six Ire1 compounds against common laboratory and corneal isolates of *A. fumigatus,* as well as the ability of the most broadly active compound, 4µ8C, to influence disease outcomes in a murine model of FK.

## 2 Materials and Methods

### 2.1 Chemical reagents used in the study

All IRE1 inhibitors were prepared in dimethyl sulfoxide (DMSO) and stored at -80 °C until use. 50 mM stock of IRE1 inhibitor III, 4µ8C (#412512 - MilliporeSigma, MA, USA), 50 mM Sunitinib (HY-10255A – MedChemExpress, NJ, USA), 10 mM Z4P (HY-153773 – MedChemExpress, NJ, USA), 10 mM STF-083010 (HY-15845 – MedChemExpress, NJ, USA), 10 mM KIRA6 (HY-19708 – MedChemExpress, NJ, USA) and 10 mM HY-114368 (HY-15845 – MedChemExpress, NJ, USA) were used. DL-Dithiothreitol (#0281 DTT-VWR, PA, USA) was prepared at 1 M concentration in sterile PBS.

### 2.2 Strains and growth conditions

The FK1 and FK2 *A. fumigatus* isolates are from fungal keratitis patients at the University of California at San Francisco-Proctor Foundation. An Af293 derivative strain that constitutively expresses the *mcherry* protein (Af293::*PgpdA-mcherry-hph*) was used throughout this study (9,19). Strains were maintained on glucose minimal media (GMM) containing 1% glucose, Clutterbuck salts, and Hutner’s trace elements, 10 mM ammonium tartrate (GMM) at pH 6.5 (20). To test the effect of the IRE1 inhibitors on the different *Aspergillus* strains, 10^6^ conidia/mL were inoculated in GMM at a concentration gradient (0 - 480 µM) and incubated at 35 °C. After 72 h, minimum inhibitory concentration (MIC) was recorded by visually inspecting the plate and looking for the growth of the fungi under the microscope. To assess the fungicidal effect of the IRE1 inhibitors, 10^6^ conidia/mL were inoculated in GMM with either 4µ8C or STF-083010 at a concentration gradient (0 -480 µM) and incubated at 35 °C. 100 µL aliquots were plated on YPD after 72 h in triplicate. These plates were incubated at 35 °C for 24 h to count the number of colony-forming units (CFU). To quantify fungal growth, the XTT metabolic assay was performed as described previously (21). XTT buffer was prepared at a stock of 0.5 mg/mL and menadione at 10 mM in 100% acetone (final conc. 1 µM). For this assay, 10^6^ conidia/mL were inoculated in GMM in 96-well plate and incubated at 35 °C for 24h to form hyphae followed by treatment with 4µ8C (0 – 240 µM) for 2h. Next, 100 µL of an XTT-menadione mixture was added to each test well and incubated further for 2 h at 35 °C. Supernatants from each well (75 µL) were assayed at 490 nm.

### 2.3 RT-PCR analyses

Fungal cultures were grown overnight at 35 °C in GMM in 6-well plates were treated with a concentration gradient of 4µ8C (0 – 120 µM) for 2 h followed by treatment with 10 mM DTT or vehicle for 2 h. RNA was then extracted from these cultures using RNeasy Mini kit (#74106 Qiagen, Germany) followed by DNase treatment using the DNase I kit (Millipore Sigma, Massachusetts, USA). The Nanodrop 2000 (Thermo Fisher Scientific, Massachusetts, USA) was used to assess the quantity and quality of the RNA. Subsequently, the RNA was standardized for cDNA conversion utilizing the ProtoScript II First Strand cDNA Synthesis Kit (New England Biolabs, Massachusetts, USA) following the provided protocol. To visualize *hacA* splicing, the coding sequencing (spanning the predicted 20 bp unconventional intron) was amplified and resolved by electrophoresis (3% agarose gel at 40 V for 10 h) as previously described (9). For qRT-PCR analysis, Luna Universal qPCR master mix (SYBR green; NEB, USA) was used on the QuantStudio 3 Real-Time PCR System (Thermo Fisher Scientific, Massachusetts, USA), and the analysis was conducted with QuantStudio Design and Analysis Software v1.5.2. Fold-expression changes were computed using the 2-ΔΔCt method, followed by analysis using MS-Excel. The primers used to amplify *hacA*, *bipA* and *pdiA* in this assay are the same as previously published (9,18).

### 2.4 Animals

<u>Inoculum</u>: 5x10^6^ conidia were incubated in 20 ml YPD at 35 °C for approximately 4 h until mostly swollen. Biomass was collected, washed 4x with PBS, and resuspended in 500 μL PBS, adjusting volumes to normalize strains to 0.8 at OD 360 nm. <u>Infections</u>: 6-8-week-old C57B6/6J (Jackson Laboratory) received intraperitoneal immunosuppression with 100 mg/kg Depo-Medrol (Zoetis, USA) on day -1. On day 0, mice were anesthetized using 100 mg/kg ketamine and 6.6 mg/kg xylazine by IP injection, and their right eyes abraded to a 1 mm diameter using Algerbrush II. Fungal inocula (5 μL) were applied to ulcerated eyes and wiped with a kim wipe after 20 minutes. The mice also received Buprenorphine SR (1 mg/kg) which was administered subcutaneously. Contralateral eyes remained uninfected per ARVO guidelines. For controls, we used algerbrushed sham-infected and untouched eyes. *<u>Micron IV</u>* slit-lamp (Phoenix Research Labs Inc., CA, USA) was used to monitor mice daily post-inoculation (p.i.) up to 72 h. This procedure was performed while the mice were anesthetized with isoflurane. *<u>Anterior segment spectral-domain optical</u> <u>coherent tomography</u>* (OCT) measured corneal thickness at 48 and 72 h p.i. using a 4x4 mm image using 12 mm telecentric lens from Leica Microsystems (IL, USA). The reference arm was set to 885 according to the manufacturer’s calibration instruction. Image analysis was conducted using the InVivoVue diver software. Corneal thickness quantification involved measuring an 11x11 spider plot covering the entire eye, and an average of 13 readings were obtained. *<u>Fluorescein</u> <u>imaging:</u>* Corneal fluorescein staining was performed to test the rate of healing of the abraded epithelium post treatment with DMSO or 4µ8C. AK-Fluor 10% fluorescein solution (Akorn, IL, USA) diluted 1:100 was applied to the corneas in isoflurane-anesthetized mice using a cotton swab and excess was wiped off and imaged using a slit-lamp Micron (Phoenix Research Labs Inc., CA, USA). *<u>Histopathological analysis</u>*: Control eyes were harvested after 72h and fixed with 10% neutral buffered formalin for 24 h followed by 70% ethanol until they were further processed. 5 μm thick sections of the eyes were stained with Hematoxylin and eosin (H&E) to evaluate host inflammatory response after treatment with DMSO or 4µ8C. *<u>Fungal burden determination:</u>* Corneas were dissected in a sterile manner, homogenized using a collagenase buffer at a concentration of 2 mg/ml, and then 100 μl aliquots were plated in triplicate on inhibitory mold agar (IMA) plates. The number of colony-forming units (CFU) per cornea was assessed after 24 hours of incubation at 35°C. *<u>Clinical scoring:</u>* Micron images were randomized and scored (0 to 4 scale) for clinical pathology by reviewers not involved with the experiment based on previously established criteria, including surface regularity, area of opacification, and density of opacification (22). *<u>Topical Treatments</u>*: For the topical treatments of sham or Af293-inoculated corneas, animals were anesthetized isoflurane and a 5 µL drop of the compound (or DMSO as a vehicle control) was applied to the ocular surface. The animals were kept under anesthesia for 10 minutes with the drop in place before being returned to their cage. The treatment schedule varied slightly across experiments (see the corresponding figure legends), but generally consisted of one treatment on the day of inoculation (4 h p.i.), three treatments (4 h apart) at 1 and 2 days p.i., and one treatment at 3 days p.i., just prior to imaging and removal of the eyes for histological or fungal burden analysis. In the case of the fluorescein experiments, the animals received three treatments 3 days p.i..

### 2.5 Statistical analyses

were performed using GraphPad Prism 10 version 10.2.0. The specific tests used for each experiment are described in the corresponding figure legends.

## 3 Results

### 3.1 Ire1 endonuclease inhibitors display antifungal activity against Aspergillus fumigatus laboratory strains and clinical isolates

We first predicted that Ire1 inhibitors would display *in vitro* antifungal activity against Af293 and potentially other *A. fumigatus* strains in which IreA influences fungal growth and or viability. We began by screening the activity of six commercially available Ire1 inhibitors ̶ including four kinase domain inhibitors (sunitinib, Z4P, KIRA6, KIRA8) and two endonuclease domain inhibitors (4μ8C, STF083010) (**Supplementary Figure 1**) in a microtiter dilution assay. Four *A. fumigatus* strains were tested in the screen, including the well-studied Af293 and CEA10 strains (both pulmonary isolates) and two uncharacterized corneal isolates from fungal keratitis patients, designated here as FK1 and FK2. No inhibitory activity was observed with any of the four kinase inhibitors even at the highest tested concentration (**Supplementary Figure 2**), suggesting that these compounds either do not adequately accumulate within the fungal cell, they do not have high affinity for the *A. fumigatus* kinase domain, or the domain itself is not essential for *A. fumigatus* growth in these isolates. By contrast, both endonuclease inhibitors fully blocked conidial germination, with the two exhibiting comparable minimal inhibitory concentrations (MICs) against the Af293 strain and 4μ8C displaying overall lower MICs against the other three (**Table 1**). To gain some insight into whether the inhibitory action of these compounds was fungistatic or fungicidal, the microtiter assay was repeated with Af293 and, following 72 h incubation, media from the wells was sub-cultured onto drug-free nutrient agar plates (**Figure 1A**). Samples corresponding to STF083010 inhibition (60-480 µM) all yielded fungal colonies, suggesting the conidia were still viable and the drug is fungistatic. All sub-culture corresponding to 4µ8C inhibition (120-480 µM), by contrast, remained sterile, suggesting the drug may be fungicidal against the conidia (**Figure 1B**). To determine if 4µ8C can similarly affect fungal hyphae (i.e. the tissue invasive form), an XTT reduction assay was performed on mycelia/biofilms cultured overnight in static culture and subsequently treated with the drug for 2 h. As shown in **Figure 1C**, the 120 and 240 µM treatments resulted in a markedly reduced or undetectable metabolic activity, respectively, altogether indicating that a similar concentration of drug is fungicidal against both conidial and hyphal form of the organism. Since 4µ8C provided the best antifungal profile, both in terms of its MIC and apparent fungicidal action, we decided to prioritize that compound for downstream characterization.

**Figure 1:**
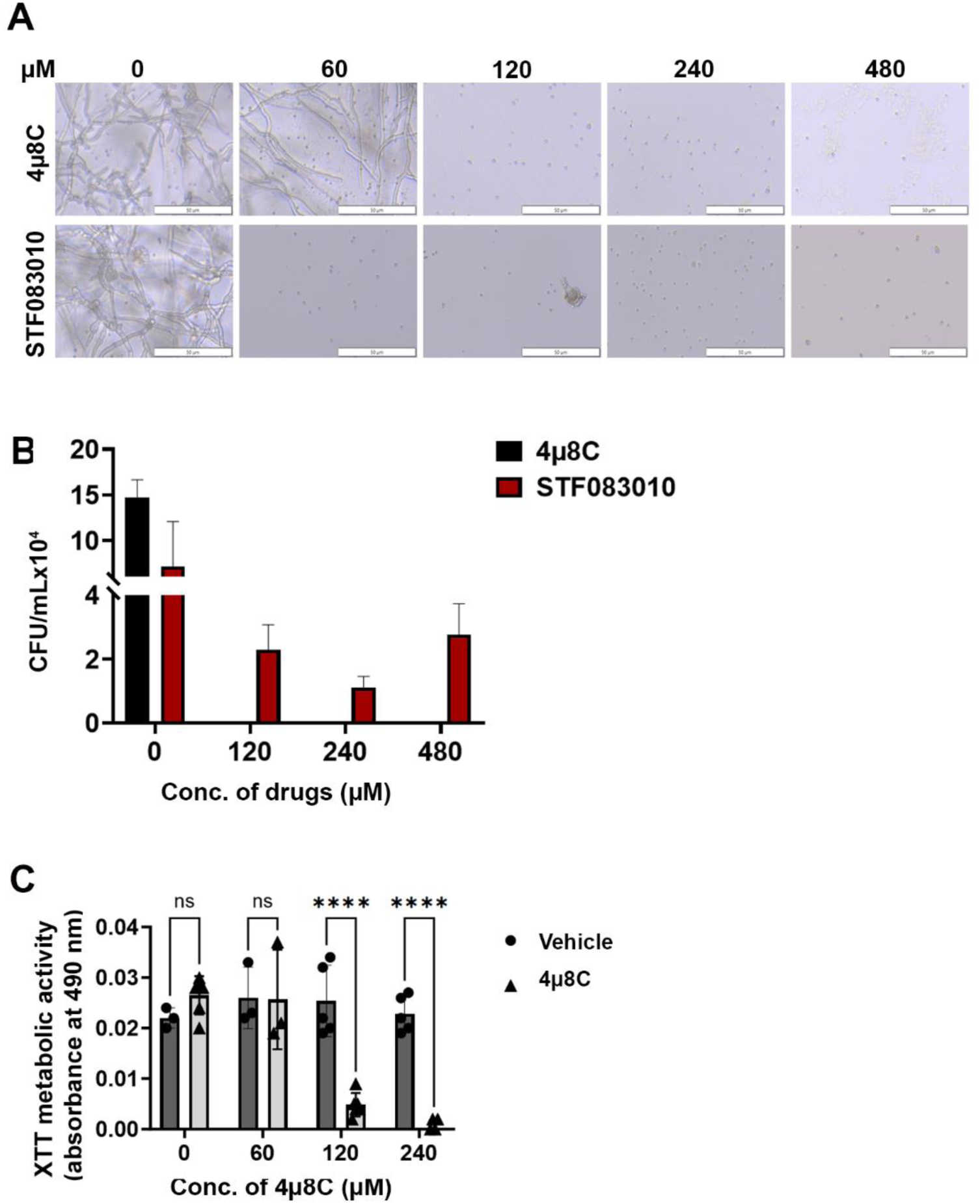
4µ8C displays antifungal activity against A. fumigatus Af293 conidia and hyphae *in vitr*o. (A) Conidia of *A. fumigatus* Af293 were inoculated into GMM broth containing the indicated concentration of 4µ8C or STF-083010 and incubated for 72 h at 35 °C. Images represent a consistent result observed across three independent experiments. (B) 100 µL aliquots were taken from wells of microbroth dilution assay, like the one shown in panel A, and spread onto YPD plates. Colonies were enumerated following 24 h incubation at 35 °C. The data reflect the mean CFU counts from triplicate wells of a single microbroth experiment. (C) Af293 biofilms generated in GMM were treated with vehicle (DMSO) or the indicated concentration of 4µ8C for 2 h at 35 °C. Metabolic activity was measured by XTT reduction. The data reflect the mean 490 nm absorbance (± SD) of triplicate wells in single experiment and groups were compared by Two-way ANOVA **** <0.001. Similar results were recapitulated across three independent experiments.

**Table 1:** Minimum inhibitory concentrations (MIC) for 4µ8C and STF 083010 against different *A. fumigatus* isolates. Data reflect a mean (± SD) of 3 separate experimental runs, except for the 4µ8C/Af293 condition which was performed across 6 runs.

|  | Af293 | CEA10 | FK1 | FK2 |
| --- | --- | --- | --- | --- |
| 4μ8C | 100 ±22.628 | 120 ±0.0 | 160 ±64.012 | 160 ±64.012 |
| STF 083010 | 80 ±32.006 | 240 ±0.0 | 240 ±0.0 | 400 ±128.024 |

### 3.2 The antifungal activity of 4µ8C corresponds to an inhibition of Ire1 activity in Af293

In biochemical assays, 4µ8C forms a Schiff base with distinct lysine residues in both the Ire1 kinase and endonuclease domains, but only inhibits the activity of the latter in treated cells (23). This lysine is conserved in the *A. fumigatus* IreA endocnuclease domain, as is the influence of the drug on *hacA* splicing in the AfS28/D141 background (18). We reasoned that if the antifungal action of 4µ8C against Af293 were attributable to an ‘on-target’ effect, then there would be a correspondence between the fungicidal MIC (ranging from 60 to 120 µM) and the inhibition of IreA function. To explore this, fungal mycelia/biofilms were developed in GMM static culture overnight, subsequently pretreated with DMSO (vehicle) or various concentrations of 4µ8C for 2 h, and finally stimulated with 10 mM DTT for an additional 2 h. Samples treated with neither 4µ8C nor DTT served as a control. The splicing status of *hacA* was then assessed by RT-PCR and gel electrophoresis as previously described (9). As expected, treatment with DTT alone resulted in an observable accumulation of the spliced/induced *hacA* product (*hacA*^i^), relative to untreated controls, and the inclusion of 30 µM 4µ8C did not have an impact on this response. By contrast, the *hacA*^i^ splice form was completely absent in both the 60 and 120 µM 4µ8C treated samples, thus establishing a putative connection between IreA function and fungal cell metabolism or viability (**Figure 2A**). Sequencing of these end-point PCR products revealed that the canonical (spliceosomal) intron at the 5’ end of the *hacA* mRNA was absent in all samples, whereas the 20 bp IreA-targeted (non-canonical) intron was absent only in the DTT-treated fungus in the absence of 4µ8C (**Figures 2C and 2D, Supplemental Figure 3**). Finally, in addition to a block in *hacA* splicing, we further observed that 60 µM blocked the DTT-mediated induction *hacA* message as well as two known HacA-regulated transcription factors, BipA and PdiA (**Figure 2B**). Taken together, these results support a model in which 4µ8C blocks UPR signaling in *A. fumigatus,* and by extension fungal metabolism and growth, through a specific action on IreA endonuclease activity, rather than a broad inhibition of mRNA processing by the spliceosome.

**Figure 2:**
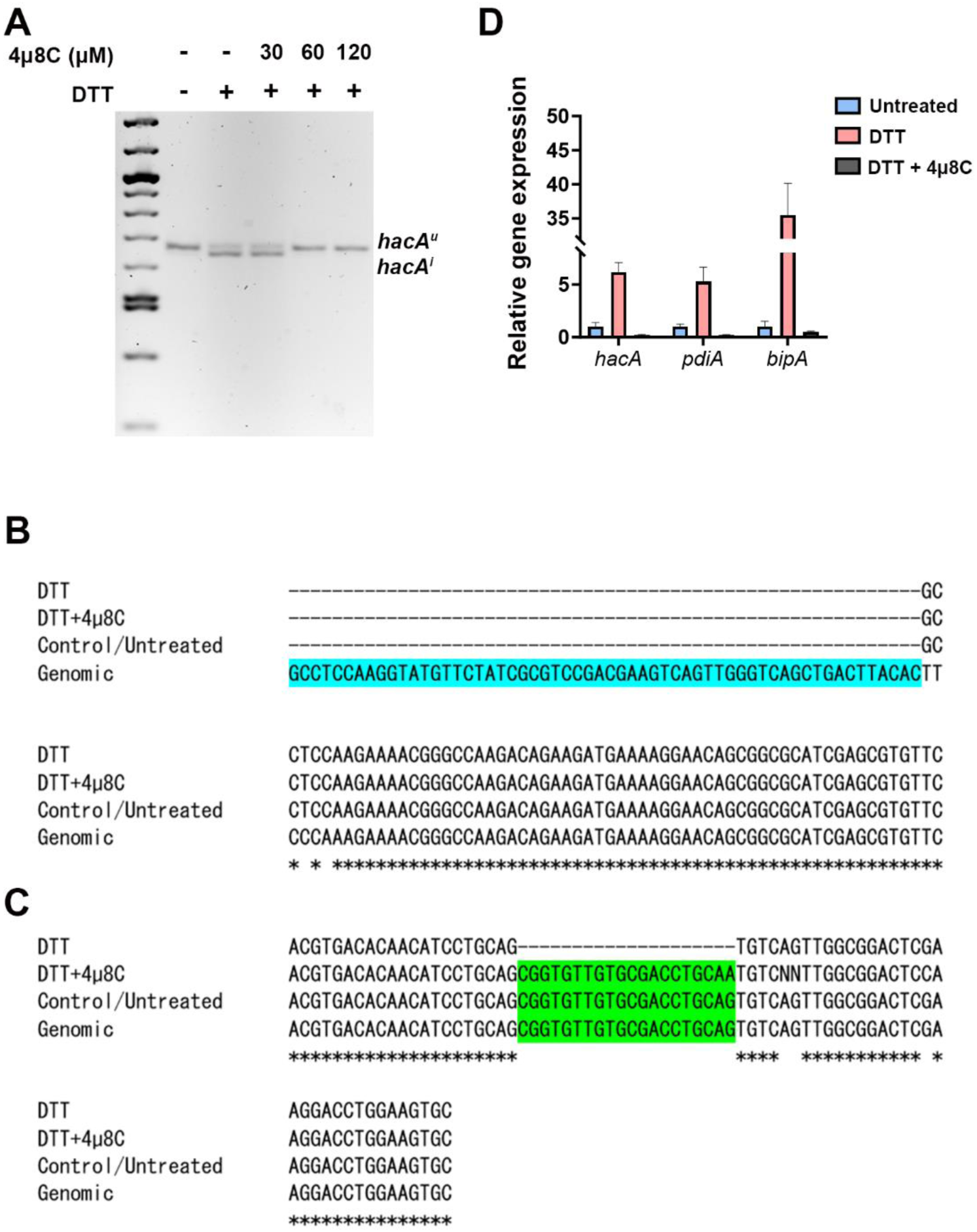
The antifungal activity of 4µ8C corresponds to an inhibition of Ire1 activity in Af293. Af293 biofilms generated overnight in GMM were pre-treated with 4µ8C or DMSO for 2 h and subsequently spiked with 10 mM DTT and incubated for an additional 2 h prior to total RNA isolation and cDNA synthesis. (A) The full-length *hacA* message was amplified and the *hacA^u^* (uninduced: 665 bp) and *hacA^i^* (induced: 645 bp) products were resolved by gel electrophoresis. The end point *hacA* PCR products were sequenced and a CLUSTAL multiple sequence alignment was performed in the regions spanning the canonical/spliceosome intron highlighted in blue (B) and the IreA-targeted non-canonical intron highlighted in green (C). (D) qPCR was performed on total *hacA* as well as two chaperone encoding genes, *bipA* and *pdiA*. The qPCR data reflect the mean 2^ΔΔCt^ calculations for triplicate fungal samples from a single experiment. Similar results were observed in an independent experiment.

### 3.3 High dose 4µ8C treatment does not impact corneal clarity but does transiently inhibit re-epithelialization

We were next interested in exploring the *in vivo* antifungal activity of 4µ8C in a murine model of FK, but first had to consider the concentration to test. Ophthalmic antifungal formulations tend to be highly concentrated to facilitate sufficient accumulation of the drug within the dense collagen matrix of the cornea, particularly in the face of rapid dilution and drainage of the drug at the ocular surface. For example, the *in vitro* MIC for natamycin against most *Aspergillus* isolates ranges between 2-32 µg/mL, but is administered at 1, 500-25,000X this concentration in the form of hourly 50 mg/mL (75 mM) drops. Similarly, voriconazole drops are often formulated at 1 mg/mL (28 mM), where the *in vitro* MIC for most *A. fumigatus* isolates is 0.25-4 µg/mL. Thus, while testing the highest concentration of 4µ8C that its solubility permits (132.24 mM) would be desirable, we had to further consider that the impact of the drug on corneal tissue homeostasis has not been explored. We therefore began by testing the ocular toxicity profiles of two relatively conservative 4µ8C concentrations, 0.5 and 2.5 mM, which correspond to approximately 5X and 40X the *in vitro* antifungal MIC, respectively.

The two drug concentrations were evaluated in sham model of FK previously developed by our group, which involves the generation of an epithelial ulcer ahead of fungal inoculation, or in this case, ulceration without the addition of fungus (9, 19). Each treatment consisted of a 5 µL drop of 4µ8C or vehicle (DMSO) applied to the corneal surface up to three times daily per the schedule described on the corresponding figure legends and methods. Corneas were tracked daily by slit-lamp imaging to assess corneal clarity and ocular surface regularity, as well as by optical coherence tomography (OCT) to measure corneal thickness and assess gross morphology. Corneas were then resected at 72 h post-ulceration for histological analysis with H&E staining. Relative to the vehicle control group, neither 4µ8C concentration had an observable impact on corneal clarity or corneal thickness, suggesting the drug did not lead to acute inflammation or edema (**Figure 3A, B** and **Supplementary Figures 4A, 4B**). Interestingly, histology revealed that whereas the epithelium of vehicle-treated corneas had reformed, those treated with either concentration of 4µ8C still appeared ulcerated (**Figure 3C** and **Supplementary Figure 4C**). To evaluate this further, we replicated the experiment and followed reepithelization (wound healing) longitudinally with fluorescein staining. Briefly, an intact corneal epithelium precludes the entry of the stain into the underlying tissue, thus giving a negative signal on fluorescence slit-lamp imaging; by contrast, ulcerated eyes give rise to diffuse yellow-green staining of the stromal layer. As opposed to the vehicle-treated eyes, which fully precluded the stain within 24 h post-ulceration, the 4µ8C-treated group remained positive for the duration of the 72 h treatment. We observed, however, that the corneas were fully healed within 3 days of stopping treatment, suggesting that the drug has only a transient impact on epithelial cell proliferation (**Figure 3D** and **Supplementary Figure 4D**). Taken together, we concluded that tested treatment regimens of 4µ8C were minimally toxic to the murine cornea and suitable for evaluation in a treatment model of FK.

**Figure 3:**
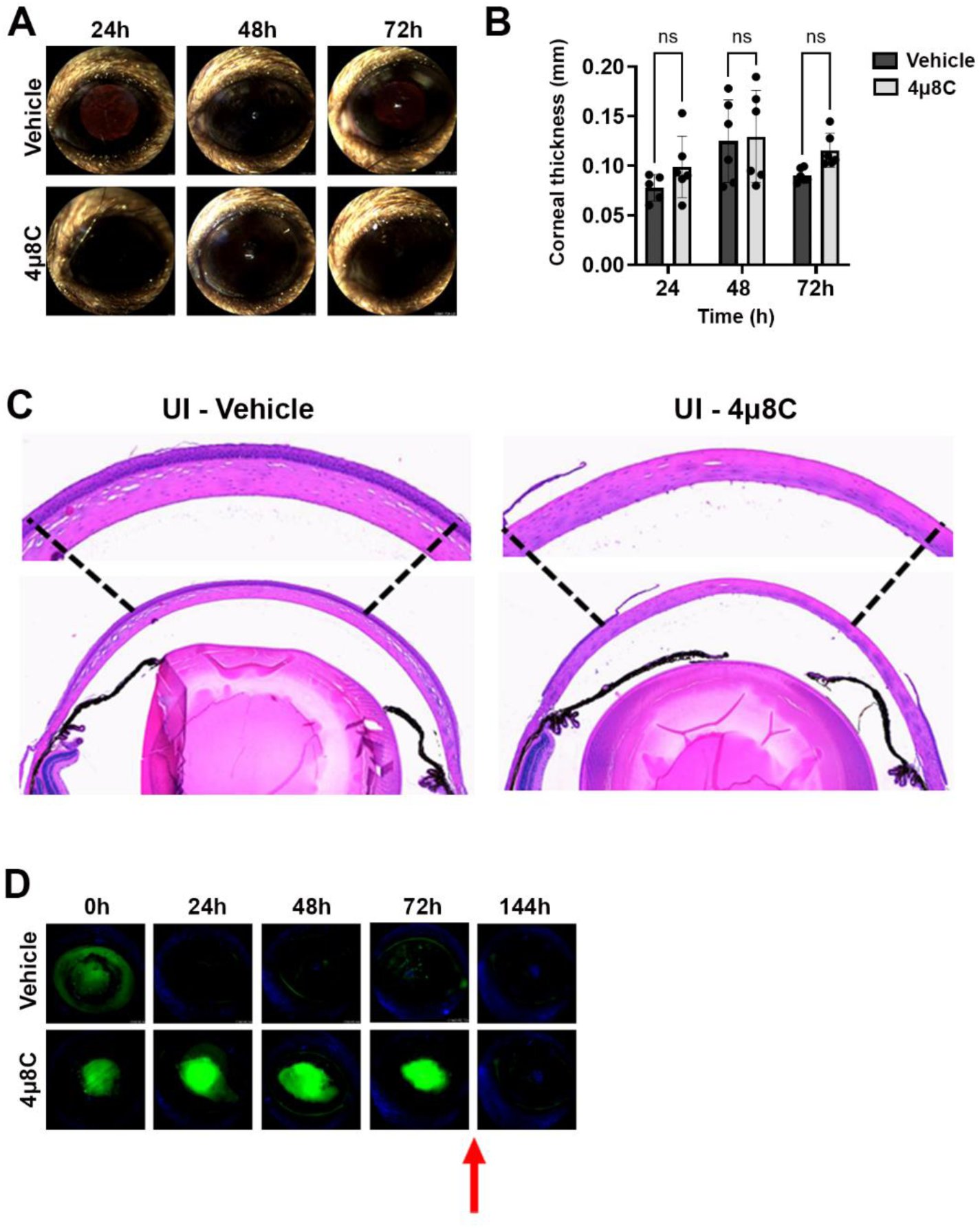
High dose 4µ8C treatment does not impact corneal clarity but does transiently inhibit re-epithelialization. Sham-inoculated (UI) corneas were treated with topical drops of 2.5 mM 4µ8C or vehicle (DMSO), once on the day on ulceration (4 h p.i.), three times (4 h apart) the following three days. (A) Representative external images taken each day post-ulceration. (B) Corneal thickness was measured daily based on OCT images (n = 6/group). Groups were compared by Two-way ANOVA; (C) Representative histological (H&E) sections taken at 72 h post-ulceration; 400X magnification. (D) In a separate experiment, ulcerated eyes were treated as described above and on each day the penetration of fluorescein was imaged by a fluorescent slit-lamp (Micron IV). The red arrow indicates that the treatment was stopped, and the eyes were monitored for an additional three days.

### 3.4 Topical treatment with 4µ8C blocks fungal growth and disease establishment in a murine model of FK

To test the impact of 4µ8C on fungal growth and disease outcomes during FK, we inoculated ulcerated corneas with swollen conidia of *A. fumigatus* Af293 (day 0) and treated on the same schedule as described above ̶ once on the day of ulceration/inoculation, three times at 24 and 48 h p.i,, and once at 72 h p.i.. Corneal disease metrics were tracked longitudinally by slit-lamp and OCT as before, but included fungal burden assessment at 72 h by homogenizing and plating corneas for colony forming unit (CFUs) measurement. We began with the 0.5 mM 4µ8C preparation, where the lack of corneal toxicity on sham/uninfected corneas observed in the previous safety trial was recapitulated. Furthermore, treatment at this concentration resulted in an approximately 50% reduction in fungal load at 3 days p.i. relative to vehicle-treated FK corneas (**Supplementary Figure 5A**). This reduction in fungal load did not, however, correspond to improved clinical disease scores or reduced corneal thickness at any of the evaluated time points (**Supplementary Figures 5B-D**), suggesting this treatment regimen is insufficient to block corneal inflammation and tissue damage associated with FK pathology. We next tested the higher (2.5 mM) concentration of 4µ8C in the same manner and observed a >90% reduction in fungal burden at 72 h p.i. that corresponded to an absence of corneal surface disease or cornea swelling at all evaluated time points (**Figure 4**). These results suggested that the higher 4µ8C dosage led to a rapid corneal sterilization that blocked the establishment of infection and downstream pathology. The relative influence of 0.5 and 2.5 mM 4µ8C on fungal load and disease development was recapitulated across multiple experiments with each concentration.

**Figure 4:**
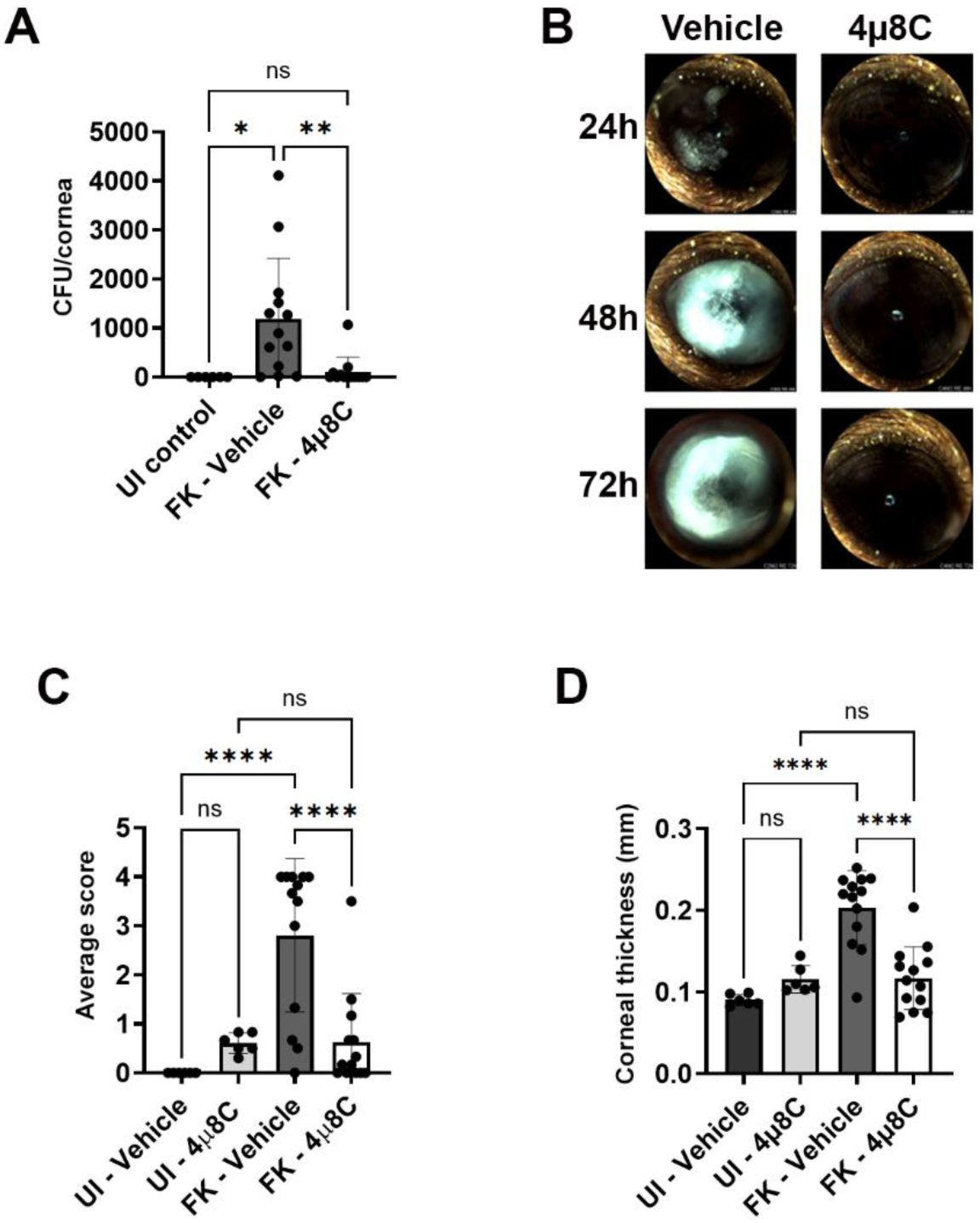
Topical treatment with 4µ8C blocks fungal growth and disease establishment in a murine model of FK. Sham (UI) or Af293-inoculated (FK) corneas were treated with 2.5 mM 4μ8C or vehicle for up to 72 h p.i. as described in the figure 3 legend. The data in all panels reflect a pool of two independent experiments (n=6 per UI group; n=13 per FK group). (A) Fungal burden at 72 h p.i.. Groups were compared by Ordinary one-way ANOVA p-value ** 0.0061, * 0.0173; (B) Representative external images taken each day p.i. (C) Average clinical scores at 72 h p.i.; Groups were compared by Ordinary one-way ANOVA p-value **** <0.0001; (D) Corneal thickness measured at 72 h p.i.. Groups were compared by Ordinary one-way ANOVA p-value **** <0.0001

## 4 Discussion

The conservation of essential proteins between fungi and mammals is an important barrier to the development of potent antifungals with acceptable host toxicity profiles. On the other hand, existing drugs developed against human proteins, particularly those in the cancer therapeutic pipeline, may display dually useful antifungal properties (24). In this report, we demonstrate that the mammalian Ire1 inhibitor 4µ8C, which blocks tumor growth in various pre-clinical models, inhibits the growth of various *A. fumigatus* isolates and can furthermore block the development of fungal keratitis when applied topically to the corneal surface (25–27). These results underscore the utility of drug repurposing efforts in the battle against medically important fungi, as well as the importance of fungal keratitis as an important experimental platform towards this end.

Our interest in developing Ire1 inhibitors as antifungals was prompted by work in *A. fumigatus* strain Af293, in which an *ireA* deletant could not be isolated and repression of the gene via a regulatable promoter halted growth under standard laboratory conditions (9). Here we demonstrate that 4µ8C fully blocks Af293 germination or hyphal metabolic activity at 60-120 µM, which aligns closely with the drug concentration that inhibits IreA endonuclease activity. Thus, the genetic and pharmacological data are seemingly in agreement and indicate that the antifungal actiivty of 4µ8C is owed to the “on-target” inhibition of an essential *A. fumigatus* protein: IreA. The possibility of “off-target” effects for 4µ8C, however, cannot be ruled out and may indeed be implicated by two lines of evidence. First, we report here that STF-083010 displays a transient inhibition of conidial germination against Af293 conidia, compared to the apparent fungicidal action of 4µ8C, despite the two compounds ostensibly targeting the same lysine residue in the IreA endonuclease domain (23). Second, work by others has established that *ireA* is not an essential gene in a D141 derivative strain (AfS28) of *A. fumigatus*, yet 4µ8C can nevertheless inhibit hyphal metabolism/viability of this strain at concentrations above which block IreA activity (8,18). Thus, while it is possible that 4µ8C inhibits other essential enzymes or metabolic processes that contribute to its *in vitro* antifungal activity, it is also likely that the exact influence of the drug is variable across *A. fumigatus* lineages/isolates. We postulate that some *A. fumigatus* lineages, such as Af293, experience a relatively higher level of ER stress during normal growth that renders IreA essential and, by extension, 4µ8C fungicidal. Alternatively, other lineages such as D141 may have recruited alternative signaling pathways that can compensate for IreA loss; in these isolates, 4µ8C would only moderately impact growth *in vitro* until higher (i.e. off-target inducing) concentrations of drug are applied. Such strain heterogeneity with respect to IreA signaling is an ongoing line of investigation in our group. Nevertheless, *ireA* is essential for the virulence of the D141 strain in the setting of invasive pulmonary aspergillosis (8), supporting its feasibilty as an antifungal target irrespective of fungal background.

We demonstrate here that topical application of 2.5 mM 4µ8C (∼40x the *in vitro* MIC in GMM) to *A. fumigatus*-infected corneas can block fungal growth and the subsequent development of disease. This effect is dose-dependent as treatment with less concentrated drops (0.5 mM) results in only a 50% reduction in fungal burden and no discernable impact on the rate of disease progression or severity. We believe these findings reflects a non-linear relationship between fungal load in the cornea and clinical pathology, such that 1) a 50% reduction in fungal antigen does not lead to a proportional reduction in the corneal inflammation and opacification or, 2) a 50% reduction in corneal inflammation is simply not clinically appreciable. Moreover, it is furthermore unclear why the 0.5 mM concentration displays incomplete antifungal activity, despite being 8X the *in vitro* MIC. We hypothesize that a combination of physical barriers, such has rapid drainage from and slow tissue pentration through the cornea, act in concert with a fungal biological response, such as drug efflux, to ultimately reduce the intracellular concentration of drug with the fungal cell. Nevertheless, the fact that an effective antifungal concentration of the drug is achievable through periodic drops alone suggest this class of compounds can be fruther optimized for ophthalmic use. Ongoing studies are aimed at elucidating the fungal physiological response to the drug as well as methods to optimize delivery to increase the effective concentration within the cornea.

We note that 4µ8C treatment did not alter corneal clarity or tissue thickness in uninfected controls, suggesting the compound does not promote corneal inflammation or edema, the latter of which can arise upon corneal endothelial dysfunction (28,29). We do observe, however, a transient block of corneal re-epithelialization, which is consistent with the known anti-proliferative effects of the compound (23,26). While at face value this represents a unwanted side-effect of the drug, we typically do not observe corneal re-peithelization of activity infected corneas in our model (9,19). In the clinical setting, we further note that a transient impact of the drug on epithelial integrity may actually be clinically useful. Indeed, a major barrier to corneal penetration of topically applied drugs is the epithelium and, consequently, this cellular layer is often debrided by clinicians to improve drug delivery. In the setting of fungal keratitis, for example, corneal debridement precedes the topical application of photosensitizers in photodynamic therapy (30,31). Thus, the antiproliferative effect of the 4µ8C may actually facilitate drug penetration in ulcerated eyes and promote its antifungal efficacy. Another important consideration is that 4µ8C also inhibits TGFβ-driven fibroblast activation and liver fribrosis *in vivo* (27,32). As myofribroblast transformation and corneal scarring are important long-term complications of FK (33–35), i.e., even if corneal sterilization is acheived, we further reason that 4µ8C treatment alone or in conjuction with other antifungals may improve visual outcomes post-infection. One final and potential impact of 4µ8C on host physiology not addressed in this study is on the ocular inflammatory response. Recently, Kumar et al. demonsrated that intravitreal injection of 4µ8C blocks the host pro-inflammatory response in a murine model of *Staphylococcus* endophthalmitis due to an Ire1-dependent activation of NK-kB signaling downstream of the Toll-like receptor 2 (38). The consequence of 4µ8C treatment in those experiments was uncontrolled bacterial growth and worsened disease outcomes. However, as fungi are also impacted by the compound, we suspect that 4µ8C may serve as a dual-edged sword that mitigates corneal damage caused by both the fungus and the host response (immunopathogenesis). We did not observe an obvious reduction in inflammation upon treatment with 0.5 mM 4µ8C, but *in vitro* studies will be required to more directly assess the impact of the compound on the pro-inflammatory response of corneal cells to fungal antigen.

In summary, the current study serves as an important proof-of-principle for the development of UPR-targeting antifungals. It also highlights the relevance of the corneal model for antifungal development more generally, as it allows for the longitudinal tracking of host toxicity and disease development before transitioning into visceral organ or systemic models of infection. Regarding FK, it is important to note that patients typically present to the clinic after the development of clinical signs of disease, including corneal haze or corneal ulcer formation. Thus, while we have demonstrated that 4µ8C can effectively kill *A. fumigatus* on the ocular surface and block disease establishment in our model, future studies will need to investigate treatment efficacy at later time points post-inoculation. We will further address the antifungal properties, both *in vitro* and *in vivo*, against other FK-relevant fungi, including *Candida albicans* and various *Fusarium* species.

## Funding and Acknowledgments

This work was supported by the National Institutes of Health (R01EY021725, P20GM134973, T32EY023202) and a Career Development Award from Research to Prevent Blindness (RPB). Core support to the OUHSC Department of Ophthalmology was further provided through P30EY021725 from NIH and an unrestricted grant from RPB. We thank Mark Dittmar and staff at the Dean McGee Eye Institute (DMEI) Animal Research Facility for animal work support, Linda Boone and Louisa Williams at the DMEI Imaging Core for histology, and the Oklahoma Medical Research Foundation Clinical Genomics Center for Sanger sequencing support. We further thank Dr. Thomas Lietman at the University of California San Francisco for the gift of the FK1 and FK2 *A. fumigatus* isolates.

---

**Figure S1:**
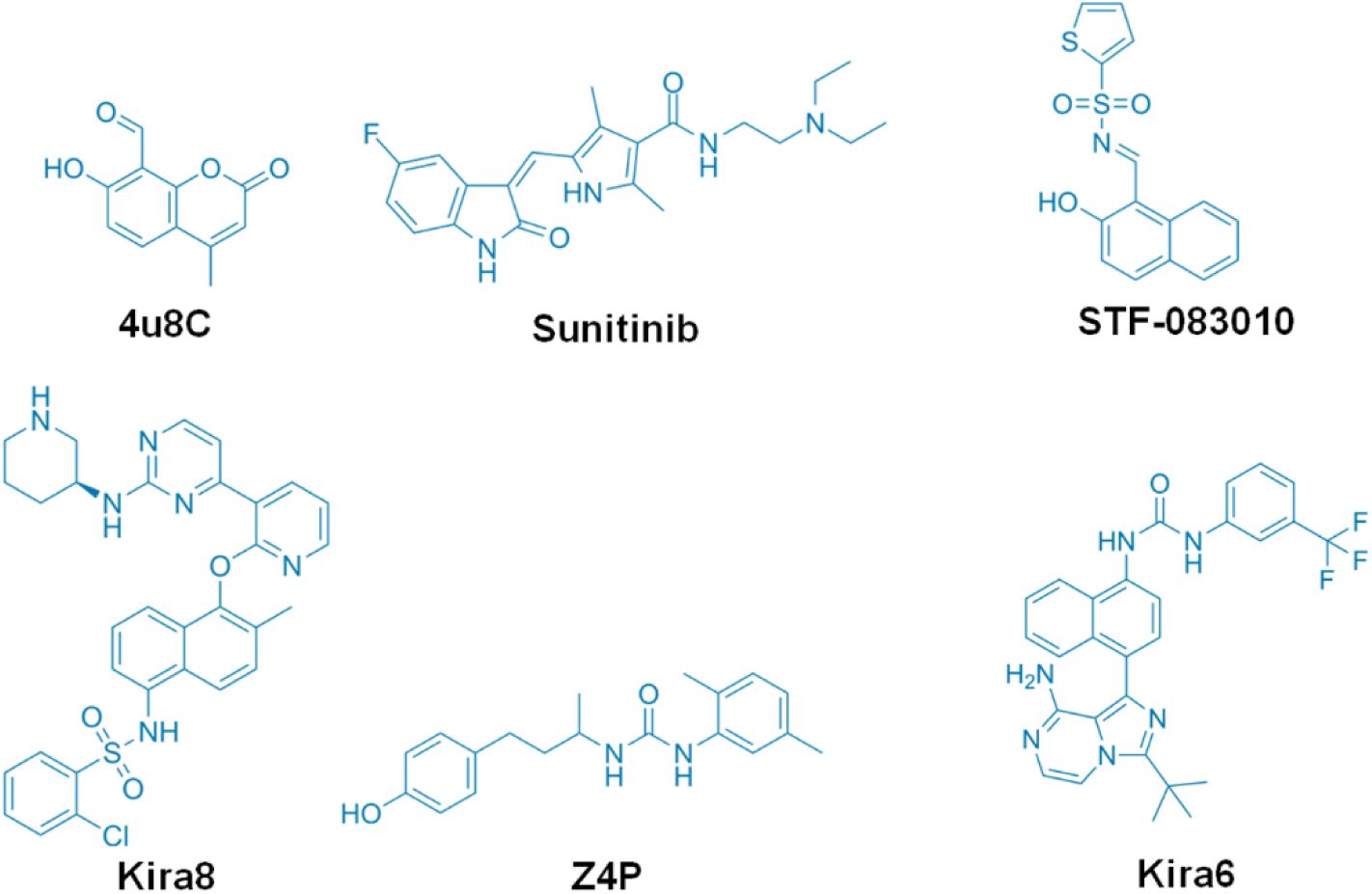
Chemical structure of the drugs screened in this study.

**Figure S2:**
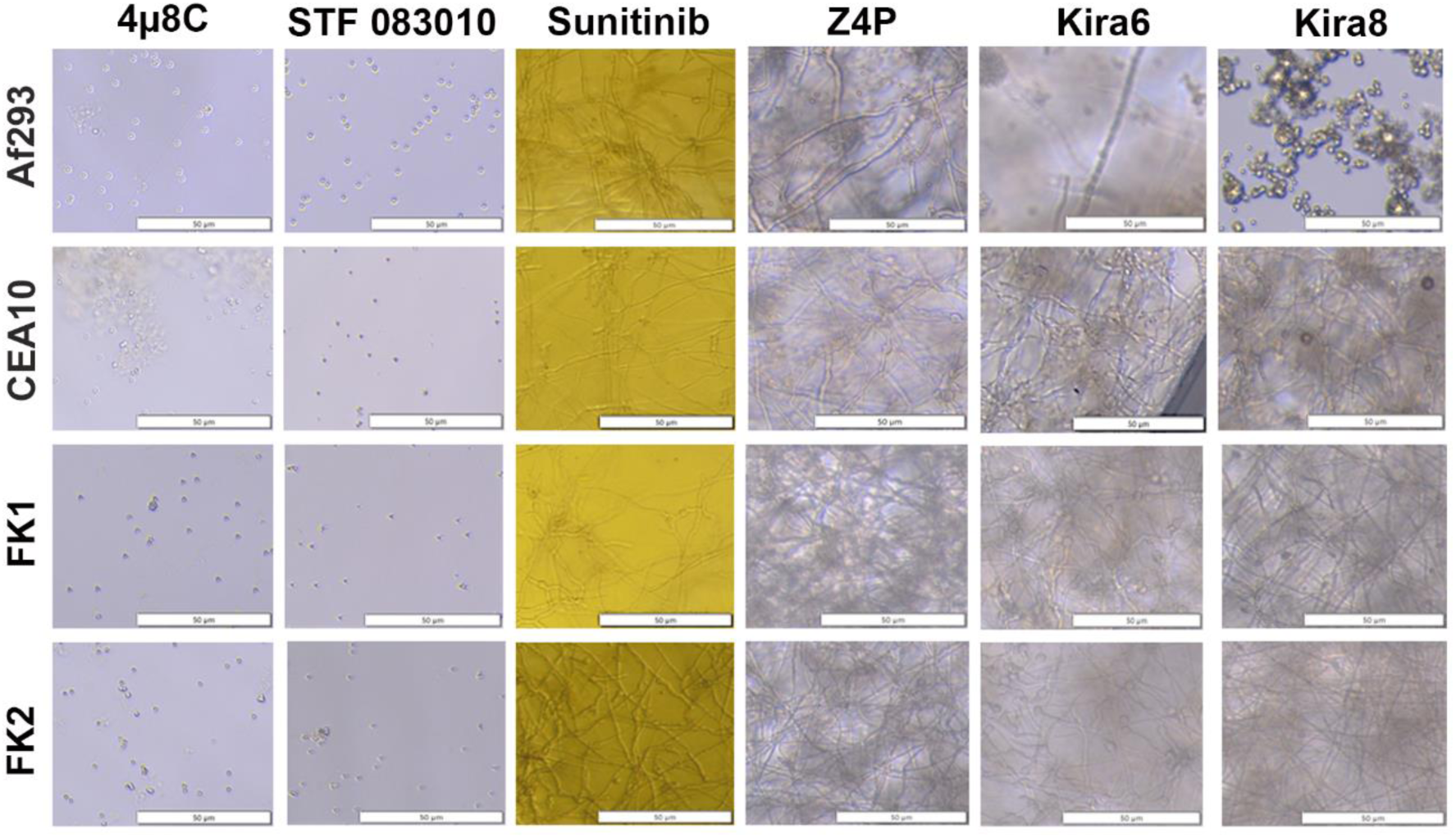
The Ire1 kinase domain inhibitors do not display antifungal activity *in vitro*. Conidia of the *A. fumigatus* isolates were inoculated into GMM broth containing the 480 µM of the indicated drug and incubated for 72 h at 35 °C. Images represent a consistent result observed across three independent experiments.

**Figure S3:**
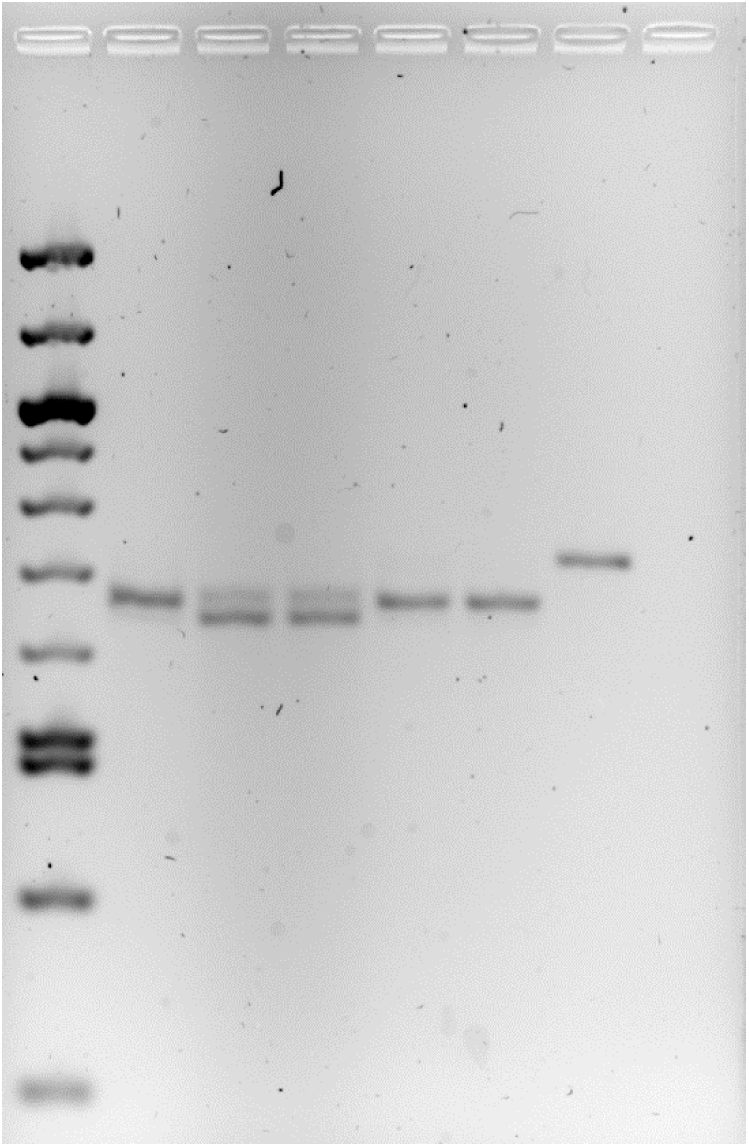
Unaltered agarose gel image corresponding to Figure 2. The original/raw ethidium bromide gel image captured from the Alliance Q9 Advanced Chemidoc (Uvitec) imaging platform. The image is unaltered in Figure 2A beyond the cropping of the rightmost two lanes, which correspond to the Af293 gDNA and no-template control amplicons, respectively.

**Figure S4:**
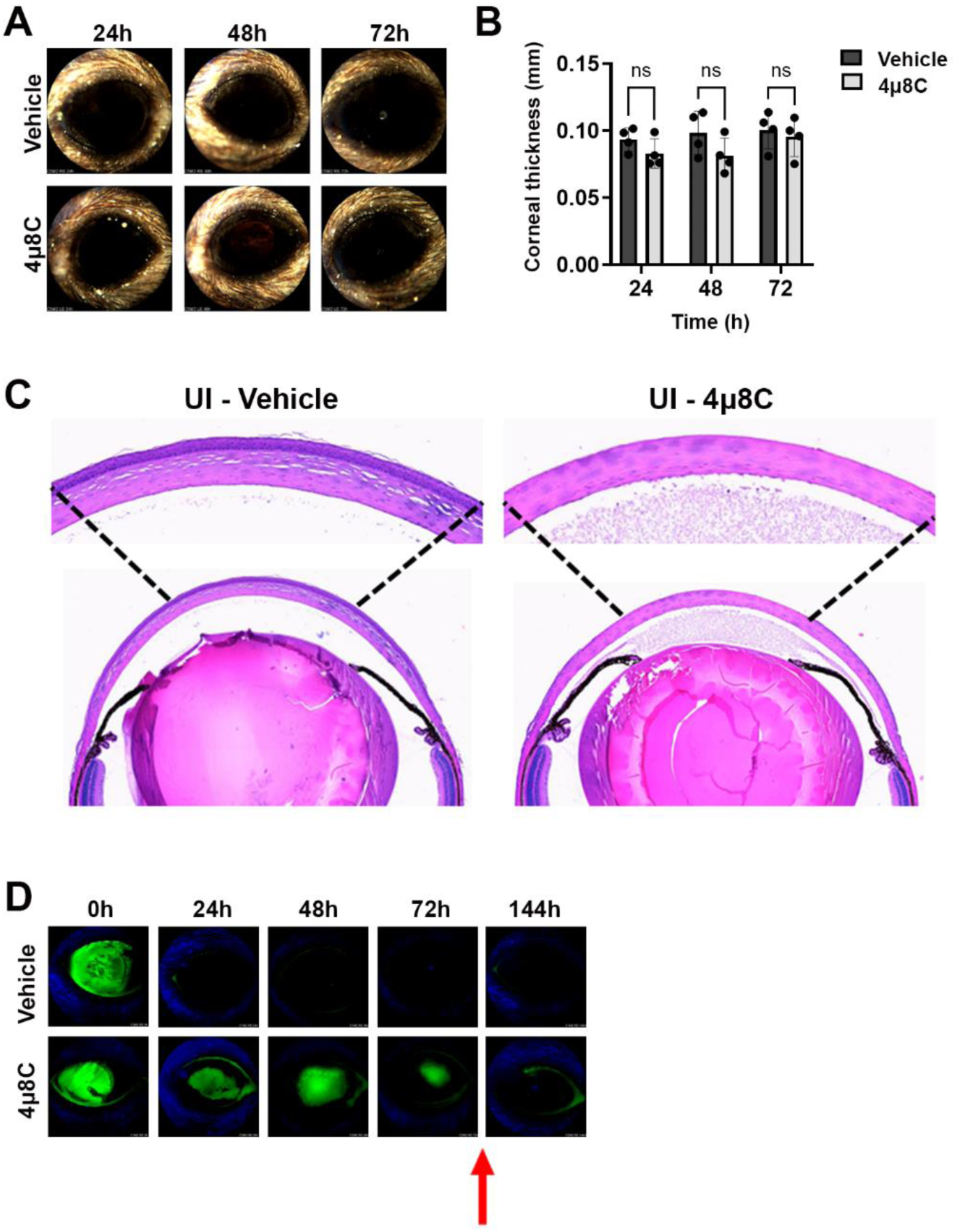
0.5 mM 4µ8C treatment does not impact corneal clarity but does transiently inhibit re-epithelialization. Sham-inoculated (UI) corneas were treated with topical drops of 0.5 mM 4µ8C or vehicle (DMSO) three times per day (4 h apart) starting on day after ulceration. (A) Representative external images taken each day post-ulceration. (B) Corneal thickness was measured daily based on OCT images. Groups were compared by Two-way ANOVA; (C) Representative histological (H&E) sections taken at 72 h post-ulceration; 400X magnification. (D) In a separate experiment, ulcerated eyes were treated as described above and on each day the penetration of fluorescein was imaged by a fluorescent slit-lamp (Micron IV). The red arrow indicates that the treatment was stopped, and the eyes were monitored for an additional three days.

**Figure S5:**
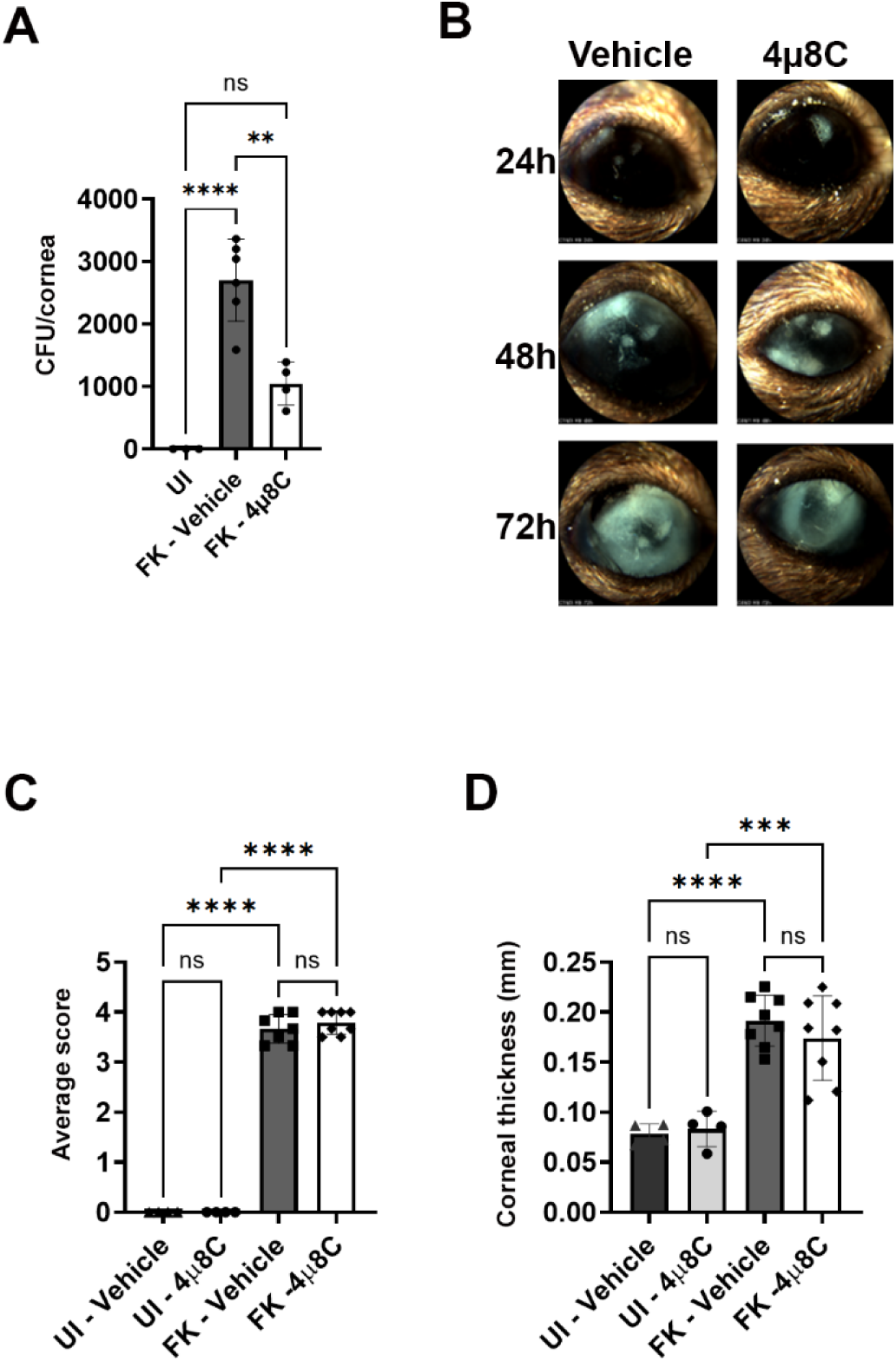
Treatment with 0.5 mM 4µ8C reduces fungal burden but has no significant impact on clinical disease severity. Af293-inoculated (FK) corneas were treated with topical drops of 0.5 mM 4µ8C or vehicle (DMSO) 3X per day starting on day after ulceration and as described in the methods. The data reflect results from a single experiment (n=4 per UI group; n=8 per FK group). (A) Fungal burden at 72 h p.i.. Groups were compared by Ordinary one-way ANOVA p-value **** <0.0001, ** 0.0012 (n=3, UI; n=6, FK-vehicle; n=4, FK-4µ8C); (B) Representative micron images over the course of the infection. (C) Average clinical scores at 72 h p.i. Groups were compared by Ordinary one-way ANOVA, p-value **** <0.0001; (C) Corneal thickness measurements at 72 h p.i.. Groups were compared by Ordinary one-way ANOVA p-value **** <0.0001, *** 0.0003.

